# Cleanifier: Contamination removal from microbial sequences using spaced seeds of a human pangenome index

**DOI:** 10.1101/2025.06.24.661305

**Authors:** Jens Zentgraf, Johanna Elena Schmitz, Sven Rahmann

## Abstract

**Motivation:** The first step when working with DNA sequence data of human-derived microbiomes is usually to remove human contamination for two reasons. First, many countries have strict privacy and data protection guidelines for human sequence data, so microbiome data containing partly human data cannot be easily further processed or published. Second, human contamination may cause problems in downstream analysis steps, such as genome assembly and binning. For large-scale metagenomics projects, fast and accurate removal of human contamination is hence critical.

**Results:** We introduce Cleanifier, a fast and memory frugal alignment-free tool for detecting and removing human contamination based on gapped *k*-mers, or spaced seeds. Cleanifier uses a pangenome index of all human gapped *k*-mers, but the creation and use of alternative references is also possible. Reads are filtered based on the gapped *k*-mers present in the index. Cleanifier supports two filtering modes: one that queries all gapped *k*-mers and one that queries only a sample of them. A comparison of Cleanifier with other state-of-the-art tools shows that our sampling mode makes Cleanifier the fastest method with comparable accuracy. Because we store the gapped *k*-mers in a probabilistic Cuckoo filter, Cleanifier has similar memory requirements to methods that use a minimizer index. At the same time, Cleanifier is more flexible, because it can use different sampling methods on the same index.

**Availability and Implementation:** The Cleanifier tool is available via gitlab (https://gitlab.com/rahmannlab/cleanifier), PyPi (https://pypi.org/project/cleanifier/) and Bioconda (https://anaconda.org/bioconda/cleanifier). The pre-computed human pangenome index is available for download at https://doi.org/10.5281/zenodo.15639519.

## Introduction

The removal of host contamination is a critical first step in metagenomics analysis projects that use human microbiomes, because of privacy concerns and to avoid complications in downstream analysis (Pereira-Marques et al., 2019), see Figure 1. For example, many public databases, such as the NCBI Sequence Read Archive (SRA) (Kodama et al., 2012), require that human-derived metagenomic sequence data is free of human reads prior to publishing. Given the ever-growing amount of microbiome data (Proctor et al., 2019), an easy-to-use, fast, and lightweight tool to remove host contamination is of high interest.

**Fig. 1:**
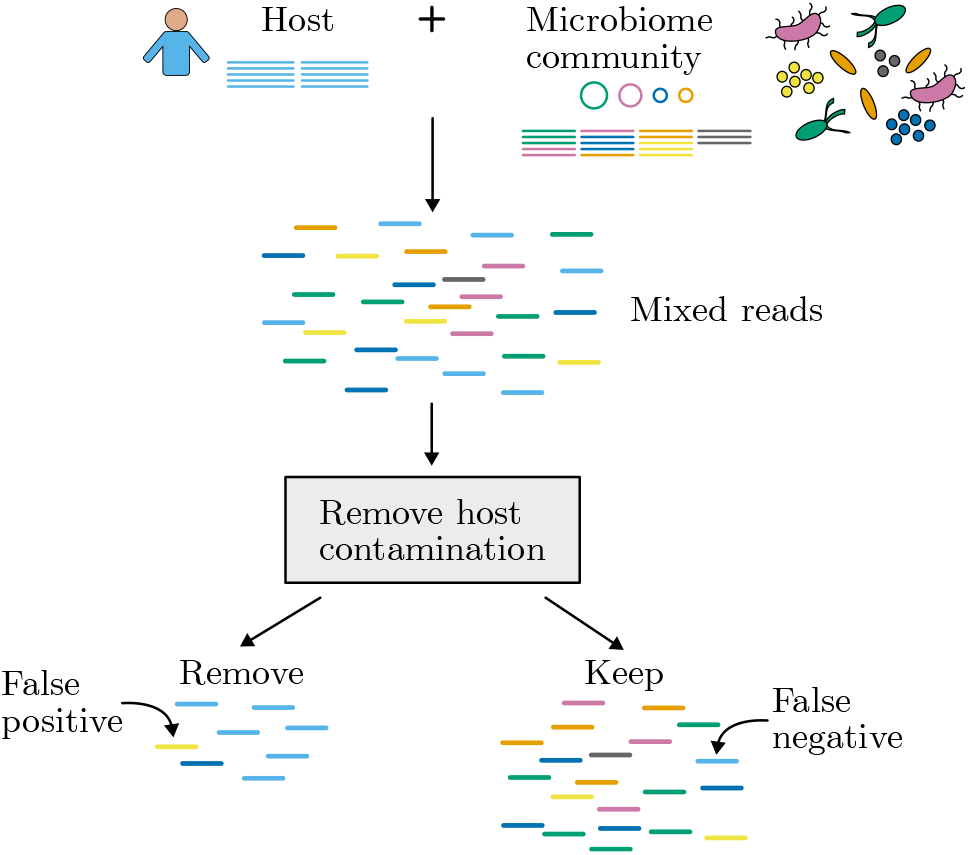
Sequencing the microbiome from a human body site, such as the gut or skin, leads to human contamination in the sequencing dataset. The goal of Cleanifier is to remove all human reads from the sequencing sample prior to publishing or downstream analysis. Wrong classification of human reads can either lead to false positives or false negatives. False positives (false removals) cause a reduced microbiome retention, while false negatives (retained human reads) lead to privacy issues.

Current tools include both alignment based and alignment free *k*-mer based methods. Table 1 shows an overview. Alignment based tools use aligners, such as Bowtie2 (Langmead and Salzberg, 2012), Minimap2 (Li, 2018) or BWA (Li and Durbin, 2009), to remove all reads that map well to the human reference. For example, KneadData (Huttenhower et al., 2020) uses Bowtie2, and Hostile (Constantinides et al., 2023) uses Bowtie2 for short reads and Minimap2 for long reads. To avoid the slow and computational expensive alignment step, *k*-mer based methods are a common alternative with a similar accuracy. The taxonomic classifiers Kraken 2 (Wood et al., 2019) and KrakenUniq (Breitwieser et al., 2018) may be used to first classify the reads and then filter out all reads that were classified as human. Other *k*-mer based tools are the Human Read Removal Tool (HRRT), which is based on the Sequence Taxonomic Analysis Tool (STAT) (Katz et al., 2021), and Deacon (Constantinides et al., 2025), which uses a human minimizer database.

**Table 1.**
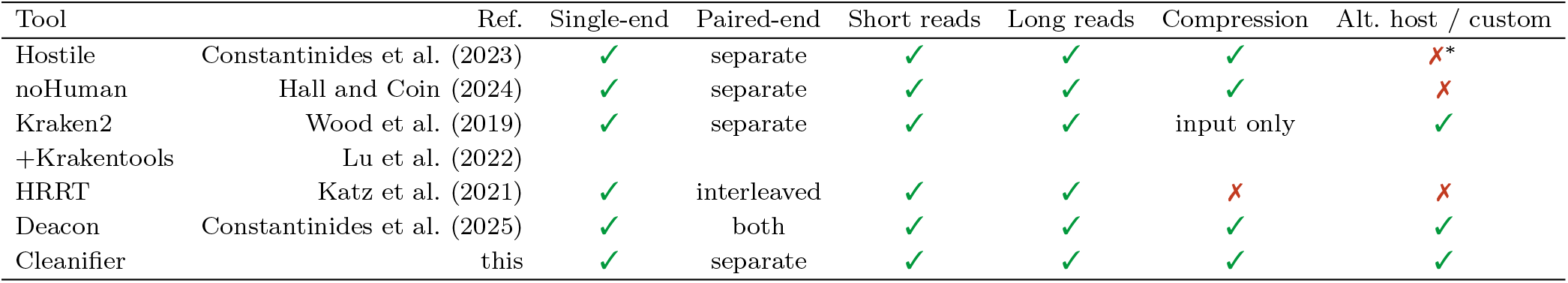
Overview of human read removal tools and supported input data. Paired-end data has to be provided either as two separate files or as interleaved paired-end format. The standard Kraken 2 database may be used to remove sequences from any host, and Cleanifier may be used to remove sequences from other hosts by building a custom host genome or pangenome database. ^∗^Hostile currently supports the removal of human or mouse reads.

We here introduce Cleanifier, a fast and accurate alignment free method that builds a database of gapped *k*-mers (spaced seeds), encompassing the human T2T reference genome (Nurk et al., 2022), all 47 genome assemblies from the Human Pangenome Reference Consortium (Liao et al., 2023), common variants from the 1000 Genome Project (Auton et al., 2015), all HLA gene variants from the IPD-IMGT/HLA database (Barker et al., 2023) and human cDNA (Dyer et al., 2025), thus containing most of the known variation in the human pangenome. The index building step only needs to be performed once per host organism and we provide the described human index online. Users only need to perform the fast filtering step using a simple command-line tool supporting single-read, paired-end, short and long read sequencing data, both in compressed and uncompressed FASTQ format.

In the Methods section, we describe the data structures used for indexing gapped *k*-mers and how we query them during the filtering step of Cleanifier. In the Results section, we compare Cleanifier with Hostile (alignment based), Kraken 2 (*k*-mer based taxonomic classifier), noHuman (Kraken 2 wrapper with custom database), HRRT (*k*-mer based) and Deacon (minimizer based). A Discussion concludes.

## Methods

### Basics

Before we describe the method in more detail, we first introduce basic definitions. We consider nucleotide sequences over the DNA or RNA alphabet Σ. Since the alphabet size is |Σ| = 4 in both cases, log_2_ |Σ| = 2 bits are needed to encode a single nucleotide.

A *k-mer* is a sequence of length *k* over Σ, i.e., an element of Σ^*k*^. For a sequence *s* ∈ Σ^∗^ with |*s*| ≥ *k* (i.e., of any length at least *k*), a substring of length *k* is called a *k-mer in s*. A *k*-mer *x* can be encoded bijectively as a 2*k*-bit integer *enc*(*x*). For *k* ≤ 32, a single 64-bit integer register or memory location suffices to store the *k*-mer.

Since DNA is double-stranded, the reverse complement of a DNA *k*-mer and the *k*-mer itself represent the same molecule and should be treated as equivalent. To represent a *k*-mer *x*, we integer-encode both *x* and its reverse complement *rc*(*x*) and then take the larger integer to represent both. Thus, the canonical code of a *k*-mer *x* is *cc*(*x*) := max{*enc*(*x*), *enc*(*rc*(*x*))}.

Example 1 *Each nucleotide is encoded as a base-4 digit by two bits (A* = (0)_4_ = (00)_2_, *C* = (1)_4_ = (01)_2_, *T* = (2)_4_ = (10)_2_ *and G* = (3)_4_ = (11)_2_*). The encoding enc*(*x*) *for a k-mer x reads x as a base-4 number*.

| $k$-mer $x$ | $enc(x)$ | $rc(x)$ | $enc(rc(x))$ | $cc(x)$ |
| --- | --- | --- | --- | --- |
| TACGC | 541 | GCGTA | 888 | 888 |
| ACGCG | 119 | CGCGT | 478 | 478 |
| CGCGT | 478 | ACGCG | 119 | 478 |
| GCGTA | 888 | TACGC | 541 | 888 |
| CGTAA | 480 | TTACG | 647 | 647 |
| GTAAG | 899 | CTTAC | 417 | 899 |

A *k*-mer is a contiguous substring of length *k*. Hence, a single variant (or sequencing error) in a read changes *k* consecutive *k*-mers. To be more tolerant against errors, previous alignment-free methods proposed to use gapped *k*-mers, also called spaced seeds, that consider *k* significant positions in a window of length *w* (Břinda et al., 2015; Wood et al., 2019; Zentgraf and Rahmann, 2022). Exactly which *k* out of the *w* positions are considered is specified by a mask defined as follows.

Given two integers *w* ≥ *k* ≥ 2, a (*k, w*)-*mask* is a string *µ* of length *w* over the alphabet {#,} that contains exactly *k* times the character # and *w* − *k* times the character . The positions marked # are called *significant*, and the positions marked are called *insignificant*, jokers, “don’t care” positions, or spaces. We call *k* the *weight* of the mask and *w* its *width* or *window length*. The pair (*k, w*) is called the *shape* of the mask. A mask *µ* may also be represented as the tuple *κ* of significant positions: *κ* = {*j* : 0 ≤ *j < w* and *µ*_*j*_ = #}.

We require that the first and last position of a mask are significant. We only consider symmetric masks to obtain the same canonical integer encodings for the forward and reverse strands. Since the robustness against substitutions depends on the positions of the significant and insignificant positions, we only consider masks with good error properties (Rahmann and Zentgraf, 2023).

Example 2 *Given the same sequence as in the previous example and the (5,7)-mask* ##_#_##, *we extract the following canonical gapped k-mer codes, where only the nucleotides in the significant positions are considered*.

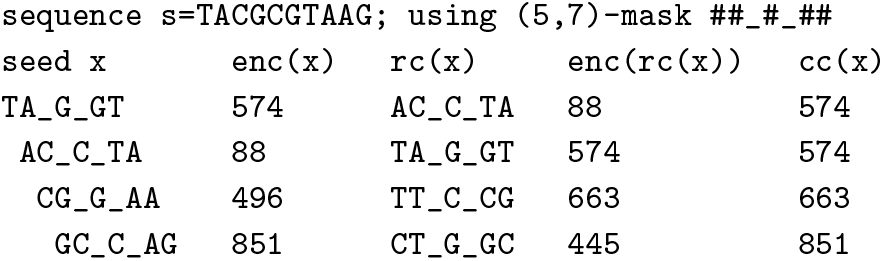

### Index data structures: Cuckoo Hash Tables and Filters

Cleanifier supports both a multiway bucketed Cuckoo hash table and a windowed Cuckoo filter for creating the gapped *k*-mer index. These variants of Cuckoo hashing have been chosen because of their low memory consumption, fast lookup times and support of online insertions, i.e., it is not necessary to know the whole key set in advance, only a good size estimate (Zentgraf and Rahmann, 2022).

### Multi-way bucketed Cuckoo hash table

Hash tables are data structures that answer membership queries for a set *S* of keys from a universe U exactly or retrieve stored information (a value) associated with each key *x* ∈ *S*. In (*d, l*) bucketed Cuckoo hashing, an array with *n* slots is divided into *b* = ⌈*n/𝓁*⌉ buckets with *𝓁* slots each. To store a key *x* ∈ U , we use *d* ≥ 2 hash functions *f*_1_, … , *f*_*d*_ : U → [*b*] := {0, 1, … , *b* − 1} to compute *d* possible bucket addresses where key *x* may be stored (together with its optional value). Hence, *x* may be stored in any of the *d𝓁* slots at *d* distinct memory locations. When checking the presence of a key, all *d𝓁* slots are searched, until either the key or a free slot is found. Since all slots of one bucket are often contained in the same cache line, we generally only have up to *d* cache misses per lookup, and for less full tables frequently only a single cache miss, yielding very fast lookup times.

The insertion of a key *x* works as follows: If we find an empty slot in one of the buckets given by *f*_1_(*x*), …, *f*_*d*_(*x*), the key is inserted in the first free slot. If no free slot is found, we remove a random key from one of the *d𝓁* slots and try to re-insert it in one of its alternative buckets. This step is repeated until either a free slot is found or a given maximum number of steps is reached. This constant threshold of steps guarantees that insertions are done in constant time, or fail. If an insertion fails, we create a larger hash table if the table is full, or we use different hash functions and redo the insertion for all elements. In the common configuration of (3, 4) bucketed Cuckoo hashing, the maximum load (fraction of non-empty slots) before insertions fail with high probability is ≈ 98%, leading to a low memory overhead (Walzer, 2023).

### Probabilistic windowed Cuckoo filter

Probabilistic filters are space efficient data structures to answer membership queries with a certain false positive probability. If we query a key *x* ∈ *S*, the filter always correctly reports the key as present. However, if we query a key *x* ∉ *S*, the filter may erroneously report the key as present with a small controllable error probability *ε*, called false positive rate (FPR). The FPR is closely linked to the space requirements of the filter, i.e., the required space increases for lower FPRs. Cuckoo filters use the same collision resolution strategy as Cuckoo hash tables and thus have the same advantages as Cuckoo hash tables, but with the additional benefit of requiring less space due to storing only a small *p*-bit fingerprint instead of an exact representation of the key *x*. Since we compare a *p*-bit fingerprint with all fingerprints stored in the *d𝓁* possible slots, the probability to falsely report a key as present is ≤ 2^−*p*^ · *d𝓁*.

The (*d, 𝓁*) windowed Cuckoo filters work similar to a bucketed Cuckoo filter, but the *n* slots are divided into *n* − *𝓁* + 1 overlapping windows instead of buckets. At each slot a new window starts and overlaps with the next *𝓁* − 1 windows. The windowed layout has been proven to admit higher load thresholds than the bucketed layout for the same (*d, 𝓁*) values (Walzer, 2023; Schmitz et al., 2025). We here use (2, 2) windowed Cuckoo filters at 95% load, resulting in 16 bits per slot for a FPR of 2^−14^ *<* 1*/*16 000, which results in approximately 16.8 bits per key.

### The indexing step

In the indexing step, using a given symmetric (*k, w*) mask, all gapped *k*-mers of the host genome are inserted into the index. More precisely, the index stores canonical codes of the gapped *k*-mers present in the provided reference sequences. The symmetry of the mask ensures that it looks identical on both the forward and the reverse strand. The index data structure is either a bucketed Cuckoo hash table (with exact canonical codes) or a windowed Cuckoo filter (with fingerprints), as described above. The advantage of the Cuckoo filter is the lower space requirement compared to the Cuckoo hash table, at the cost of a small FPR, which we show to be negligible in practice (see Results).

### Indexing the human pangenome

We provide a precomputed pangenomic index for filtering human reads using the following (29, 33) symmetric gapped *k*-mer mask ‘######_#######_###_#######_######’. It guarantees at least 50 out of 100 covered positions for every substring of length 100 of the read for which there exists a substring of the genome that differs by at most 3 substitutions (Rahmann and Zentgraf, 2023). The index is based on the T2T reference genome (Nurk et al., 2022), which provides the benefit that it does not contain undefined bases. By using gapped *k*-mers, the approach is already robust against SNPs, which is further improved by including the variants from the 1000 Genome Project (Auton et al., 2015). For this, we analyze the VCF files based on the T2T-reference for all 3202 genomes, extract all variants, namely substitutions, insertions and deletions, with an allele frequency of at least 1%. We replace the region in the reference with the provided alternative sequence and add all gapped *k*-mers ±50 bp of the new sequence to the index. In addition, we include the gapped *k*-mers from all 47 pangenome assemblies of the Human Pangenome Reference Consortium (Liao et al., 2023). Since the MHC is a highly variable region in the genome, we further add all gapped *k*-mers from HLA gene variants stored in the IPD-IMGT/HLA database (Barker et al., 2023). To be able to remove mRNA contamination, we additionally add all gapped *k*-mers from the human cDNA sequences derived from Ensembl genes (Dyer et al., 2025).

A comparison concerning classification accuracy between this pangenomic index and an index that only contains the gapped *k*-mers of the T2T reference genome is provided in the Supplement.

### Filtering microbiome samples

To decide whether a read *s* belongs to the host organism or not, we query the gapped *k*-mers of *s* in the human index. If we find a gapped *k*-mer in our index, we label the base pairs overlapping with this gapped *k*-mer as *covered*. Given a threshold *T* ∈ [0, 1], a read is then classified as originating from the host genome if at least *T* |*s*| basepairs are covered.

Cleanifier supports two classification modes, a faster sampling mode and a slower sensitive mode:

- In sensitive mode, all gapped *k*-mers of the read are queried. If a gapped *k*-mer exists in the index, the significant positions in the read are marked as covered.
- In sampling mode, for a mask of shape (*k, w*), we query the first gapped *k*-mer of the read and then skip the next ⌊*w/*2⌋ consecutive gapped *k*-mers before we query the next one, and so on. Hence, every base pair has a chance of being covered by two distinct gapped *k*-mers. If a gapped *k*-mer is present in the index, we count all *w* positions (not only the significant ones) as covered.

The above holds for single-end reads. To process paired-end reads, each read of a pair is first classified independently. If at least one of the two ends is classified as a host read, the whole pair is removed.

### Implementation details

Cleanifier is implemented as a workflow-friendly command-line tool in Python using the `numba` package (Lam et al., 2015) for just-in-time compiling the Python code. We use our own implementations of windowed Cuckoo filters (Schmitz et al., 2025) and bucketed Cuckoo hash tables (Zentgraf and Rahmann, 2022) that use double quotienting for storing partial keys (without sacrificing exactness) to reduce the space requirements of the hash table.

### Parallelization

Our implementation parallelizes both the indexing and the filtering step of Cleanifier using a consumer-producer architecture. For indexing, parallelization is over sub-tables, with a single thread being responsible for inserting keys or fingerprints into a sub-table. An additional outer hash function selects the sub-table for each *k*-mer and delegates insertion to the corresponding inserter thread. For filtering with its read-only workload, parallelization is trivial over chunks of reads or read pairs. The threads communicate over lock-free buffers that use atomic volatile `ready_to_read` and `ready_to_write` flags.

### I/O and consumer-producer architecture

For indexing, a dedicated I/O thread reads (chunks of) the genomic input sequences into buffers that are consumed by threads that split the reads into their gapped *k*-mers and writes their canonical codes into other buffers. Inserter threads (one per sub-table) consume theses buffers and perform the insertions.

For filtering, a dedicated I/O thread reads the FASTQ files into multiple buffers. Compressed input and output is supported via external tools in separate processes with data being transferred via pipes. For paired-end data, each input buffer is divided into two halves, the first half contains the reads of the first pair and the second half of the other pair. The classifier threads consume the buffers from the I/O threads, classify the reads by querying the gapped *k*-mers and write the classification results into their respective output buffers. A writer thread consumes the output buffers from the classifier threads and writes the reads that are not classified as human to the output FASTQ files.

### Index in shared memory

Cleanifier provides the option to load the index into shared memory once where multiple independent Cleanifier instances may use it simultaneously for (read-only) filtering. This has two advantages in many-sample scenarios. First, this saves the time to load the index into memory each time a Cleanifier process is started. Second, this reduces the required memory and enables parallelization over multiple instances without an increasing memory footprint.

## Results

First, we describe the human and microbiome data and the tools used in our evaluation. We then compare all tools with respect to their filtering accuracy, running time and memory requirements on short and long read sequencing data.

Benchmarks were run on an Ubuntu (24.04 LTS) server with two AMD EPYC 9534 64-Core Processors, 1.5TB of 4800-MHz DDR5 memory and a KIOXIA CD8P SSD. We measured running times and maximum memory usage with `/usr/bin/time -v`.

### Datasets

We used human and microbiome datasets as detailed below. Dataset URLs are provided in the Supplement.

#### Human data

We used human whole genome sequencing datasets provided by the Genome in a Bottle (GIAB) consortium (Zook et al., 2016), HG002–HG006. In principle, these datasets should contain only human reads, but some contamination with viruses or bacteria or technical artifacts cannot be excluded. It is therefore unclear whether we should expect that 100% of all reads should be removed. We evaluated the performance for short reads on paired-end Illumina data, downsampled to contain 50 million 2×125 bp read pairs from each dataset HG002–HG006. For long reads, we used Pacific Biosciences (PacBio) reads of HG002, downsampled to 5 million reads.

#### Microbiome data

We downloaded the CAMI 2 challenge human microbiome dataset (Meyer et al., 2022; Fritz et al., 2021) for 5 different body sites (gastrointestinal, airways, oral, skin and urogenital) as a gold standard that contains only microbial reads. The dataset is a mixture of reads from different species based on the microbiome profiles from the Human Microbiome Project (Huttenhower et al., 2012). We combined the available FASTQ files, such that each file per body site contains 50 million paired-end Illumina short reads (2×150 bp) and 5 million PacBio long reads, respectively.

### Tools

We evaluated Cleanifier against five state-of-the-art methods for human read removal: Hostile, Kraken 2 with Krakentools, noHuman (based on Kraken 2), HRRT and the recent Deacon. We ran all tools using 8 threads and their respective default parameters, unless explicitly stated otherwise. We provide a brief description of the evaluated tools below.

We used Cleanifier with the human pangenomic index described above and a coverage threshold of 0.5 (the default) for classification. We ran four different Cleanifier versions, combining either the exact hash table or the probabilistic filter with an FPR of 2^−14^ with the sensitive mode or the sampling mode for classification.

Hostile (Constantinides et al., 2023) is an alignment-based approach to remove human contamination by aligning all reads to the human genome using Bowtie2 for short reads and Minimap2 for long reads. We ran Hostile with its default index comprised of the human T2T reference (Nurk et al., 2022) and HLA variants from the IPD-IMGT/HLA database (Barker et al., 2023).

Kraken 2 (Wood et al., 2019) is a fast *k*-mer based taxonomic classification tool for metagenomic sequencing data. We evaluated Kraken 2 using the standard database (available for download at https://benlangmead.github.io/aws-indexes/k2). After taxonomic classification, reads that are not classified as *Homo sapiens* (taxonomy ID: 9606) are extracted using Krakentools (Lu et al., 2022).

noHuman (Hall and Coin, 2024) is a Kraken 2 wrapper specifically built to remove human contamination. It uses a custom Kraken database built from all genomes available in the Human Pangenome Reference Consortium (Liao et al., 2023).

HRRT is part of the SRA Taxonamy Analysis Tool STAT (Katz et al., 2021). Reads are removed by querying *k*-mers in a MinHash-based index that contains all *k*-mers present in human derived eukaryotic species; excluding all *k*-mers that are known to also occur in non-eukaryotic species.

Deacon (Constantinides et al., 2025) builds a pangenomic index that contains all human minimizers using a fast SIMD minimizer construction (Koerkamp and Martayan, 2025). Reads are classified as human if at least 2 minimizers are found in the index. We use the prebuilt `panhuman-1` index (available at https://github.com/bede/deacon).

### Results on short reads

We report the fraction of correctly classified reads (i.e., correctly removed reads from the human datasets, and correctly retained reads in the microbial datasets) in Figure 2. We note that no tool removes more than 98% of the human reads. This does not necessarily mean that all tools miss at least 2% of the reads. All of these 5 whole genome sequencing datasets (HG002–HG006) may contain non-human contamination (e.g. Epstein-Barr virus, phiX phage), or technical artifacts. The correct set of reads to be removed is unknown, and probably not 100%, but we still may assume that more removed human reads are better. On the other hand, 100% of retained microbial reads are certainly desirable, and almost all tools achieve close to 100% retention rate, with the lowest value of any tool on any dataaet being 99.983%.

**Fig. 2:**
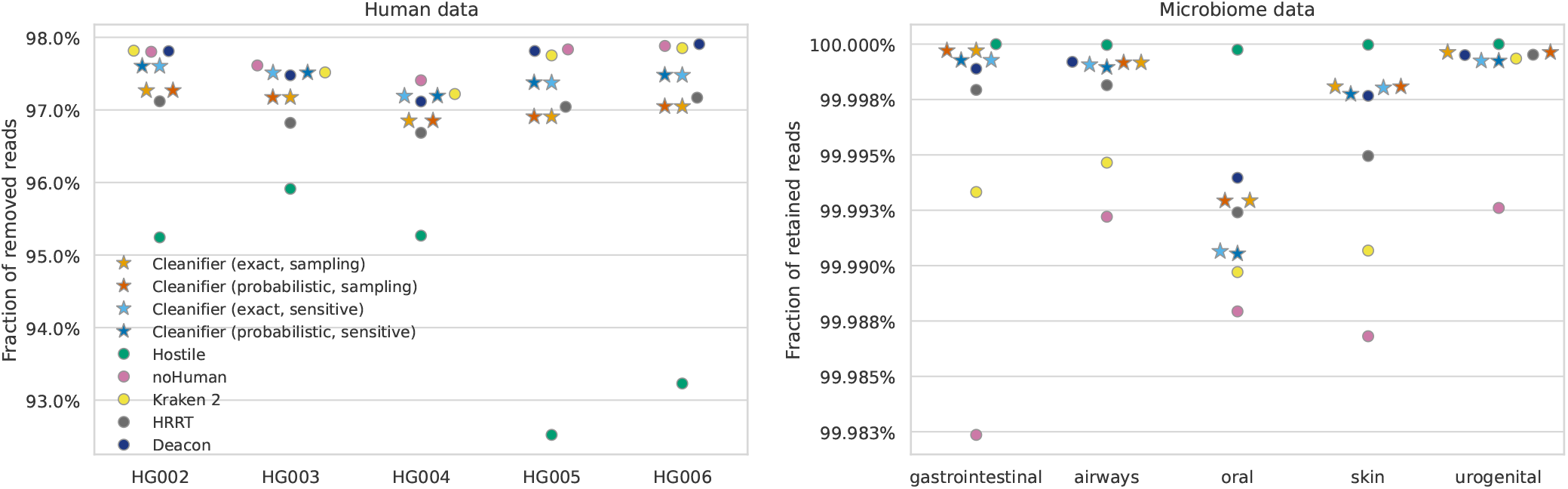
Results on Illumina paired-end short reads for human datasets (left panel; fraction of removed reads) and microbiome data (right panel; fraction of retained reads); higher is better. The different Cleanifier versions are indicated by differently colored stars; other tools by circles.

Kraken 2 and noHuman are sensitive to human reads, resulting in the highest removal rate on human data, but also on the lowest retention rate on microbiome data.

In contrast, Hostile has the best microbiome retention among the tools, but it is much less sensitive to human reads, to a degree where it may be considered unsafe to use in privacy-sensitive settings.

All Cleanifier versions, Deacon and HRRT have a high accuracy on both human and microbiome data, with Cleanifier having a slightly higher accuracy compared to HRRT and Deacon performing slightly better on human sequence removal.

Among the Cleanifier versions, the exact and probabilistic data structures have almost the same accuracy on all samples, showing that for the default coverage threshold of 0.5, the small FPR incurred by the filter for a single *k*-mer query is negligible in practice, as several *k*-mer matches are required to classify a read as human. The sampling mode (red/orange stars) of Cleanifier is slightly less sensitive to human reads compared to the sensitive mode (blue/cyan stars): On average, 0.38% of all reads are additionally retained in the human datasets. On microbiome data, the sampling and sensitive mode have a comparable accuracy.

To investigate the question which reads from the human datasets may be in fact of non-human origin, we created Venn diagrams for the retained human reads for Cleanifier (probabilistic) in both sampling and sensitive mode and each other tool, based on HG002 (Figure 4). More detailed comparisons can be found in the Supplement. The reads classified as non-human by Cleanifier (in both sampling and sensitive mode) are almost a true subset of the reads retained by Hostile and share a high overlap with noHuman, Kraken, HRRT, and Deacon, suggesting that those reads may indeed be contaminants or other technical artifacts.

**Fig. 3:**
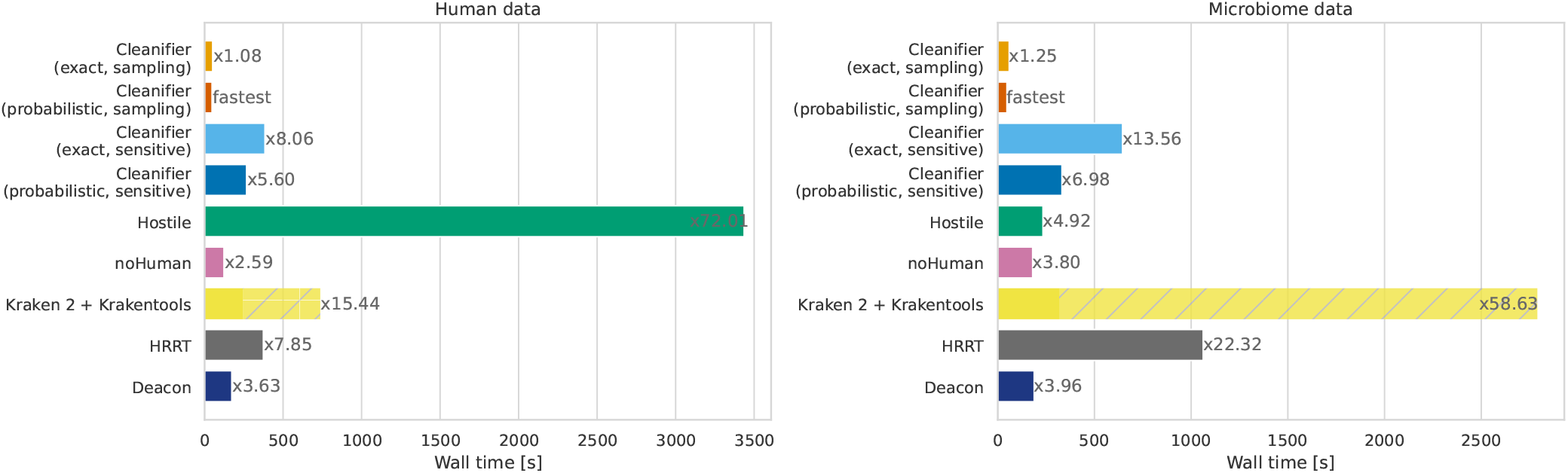
Running time in seconds on Illumina short read datasets with 50 million paired-end reads (left panel: human data; right panel: microbiome data). All tools are run with 8 threads. The running time is averaged over 5 human and microbiome datasets, respectively. Factors next to bars show running times relative to the fastest tool (Cleanifier in sampling mode using a probabilistic Cuckoo filter).

**Fig. 4:**
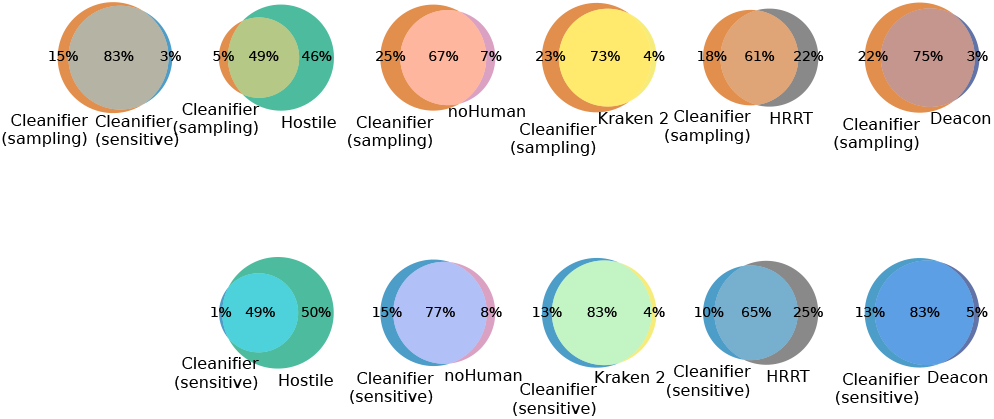
Overlap between retained human reads of HG002 between Cleanifier (probabilistic) and the other tools.

### Running time on short reads

We measured wall clock running times for reading, classifying and writing uncompressed FASTQ files and configured all tools to output only the microbial reads. Figure 3 shows the wall times for FASTQ files with 50 million paired-end short reads using 8 threads, shown separately for human and microbial reads. Speedup factors for using Cleanifier with different numbers of threads can be found in the Supplement.

On both human and microbiome data, Cleanifier (sampling) is fastest, while the sensitive mode is slower than both Cleanifier (sampling) and noHuman, which is the second fastest tool, almost on par with Deacon. The next fastest tools overall are Cleanifier (probabilistic, sensitive), HRRT and Cleanifier (exact, sensitive). All of them are relatively slower on microbiome data than on human data, which can be explained by the fact that all *k*-mer lookups fail in the hash table. HRRT suffers more from this than the two sensitive Cleanifier versions.

Taxonomic classification with Kraken 2 using the standard database it slower compared to the smaller database used in noHuman. The overall running time of Kraken 2 + Krakentools is significantly slower, due to the slow filtering with Krakentools. In particular, Krakentools is very slow if large FASTQ files have to be written as is the case for the microbiome data. In the case of human data, almost all data is discarded, resulting in a lower overhead of Krakentools.

As expected, Hostile is slowest on human data (72 times slower than the fastest method) due to the slow alignment step. Surprisingly, it is among the fastest tools on microbiome data. The reason is that no alignment needs to be constructed for most of the reads since no start seed (short exact match) is found to initiate an alignment. In practice, human contamination in microbiome samples can reach up to 90% (Gao et al., 2025), requiring tools that are fast for classifying both human and non-human reads.

### Results on long reads

We also measured the accuracy on long reads (Figure 5). All tools, except Kraken 2 and noHuman, have an accuracy of almost 100% on the microbiome datasets. On the human HG002 dataset, the accuracy of the Cleanifier variants and that of Hostile is close to 100%. (Note that this is a completely different dataset than the HG002 short read dataset, with a very different library preparation and likely without contamination due to filtering by read length.) The accuracy of all tools on the human dataset is > 99.9%, with that of Deacon being below the Cleanifier variants but above Kraken 2, HRRT and Hostile.

**Fig. 5:**
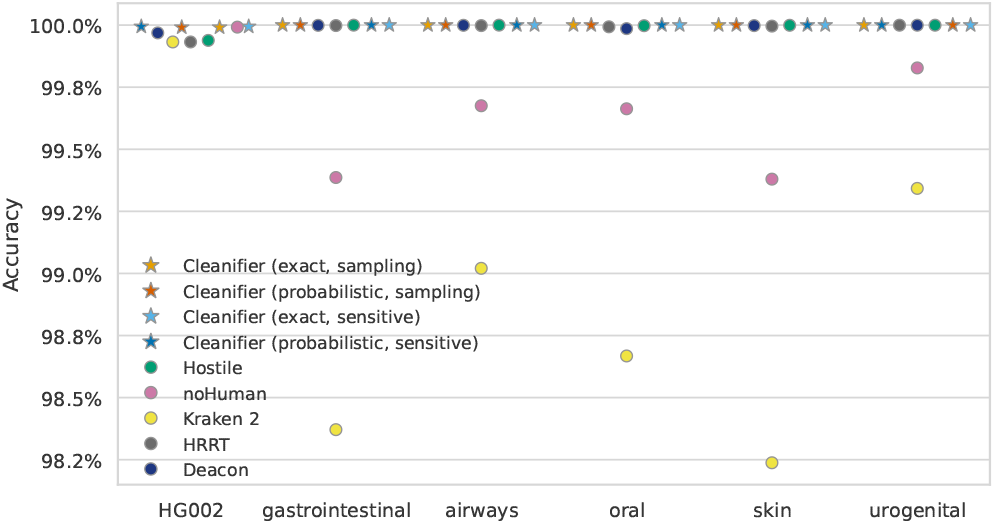
Accuracy, i.e., fraction of removed reads (human dataset HG002) or fraction of retained reads (microbiome datasets) for PacBio long reads; higher is better.

**Fig. 6:**
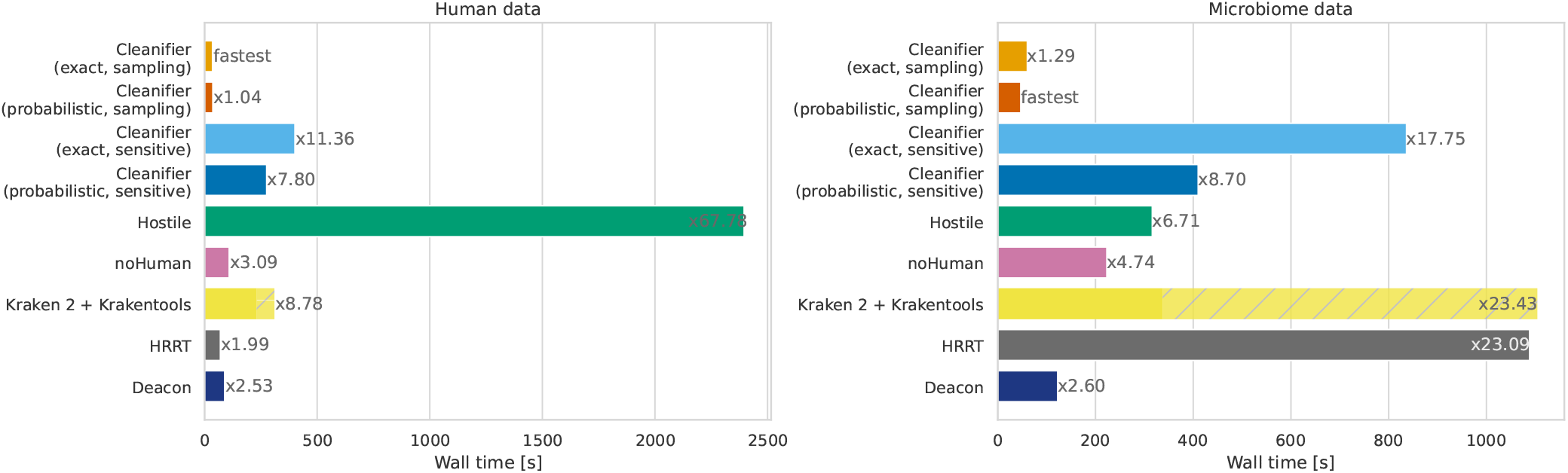
Wall clock running time in seconds for filtering 5 million long reads with 8 threads (left panel: human data; right panel: microbiome data). The running time for microbiome data is averaged over the 5 samples. Factors next to bars show running times relative to the fastest tool.

### Running time on long reads

The relative wall clock running time between the tools on long reads is similar to the time on short reads. The sampling versions of Cleanifier are fastest on both types of datasets. On human data, the running times of HRRT, Deacon and noHuman are slightly slower (2–3 times), the sensitive mode of Cleanifier and Kraken 2 are significantly slower (8–12 times), and Hostile is extremely slow (68 times slower) because of the expensive alignments. On microbiome data, again Deacon and noHuman are slightly slower than the fastest tool (2.5 to 5 times), whereas the running time of HRRT, Kraken 2, and the sensitive exact version of Cleanifier increases sharply (18 to 24 times). In contrast, the running time of Hostile decreases and becomes comparable the the sensitive mode of Cleanifier (probabilistic). In summary, the sampling versions of Cleanifier, Deacon, and noHuman show consistently fast running times in all situations.

### Memory requirements

Table 2 shows the memory requirements measured for filtering short read data. The memory is mostly used for holding the index data structure in memory, and to a lesser degree for sequence buffers. Hostile, HRRT and Deacon have the smallest memory footprints below 5 GB. Between 5 GB and 7 GB is achieved by noHuman and Cleanifier (using the probabilistic Cuckoo filter with an FPR of 2^−14^). The exact representation of Cleanifier requires still under 16 GB, whereas the comprehensive Kraken 2 database needs much more memory, close to 90 GB. However, as seen in the case of noHuman, a smaller Kraken 2 database suffices to get a high accuracy.

**Table 2.** Maximum memory usage in GB; measured on short read samples.

|  | Cleanifier |  | Hostile | noHuman | Kraken 2 | HRRT | Deacon |
| --- | --- | --- | --- | --- | --- | --- | --- |
|  | exact | prob. |  |  |  |  |  |
| Human | 13.85 | 6.90 | 3.86 | 5.22 | 88.83 | 4.37 | 4.72 |
| Microbiome | 13.85 | 6.90 | 3.65 | 5.38 | 90.46 | 1.00 | 4.73 |

**Table 3.**
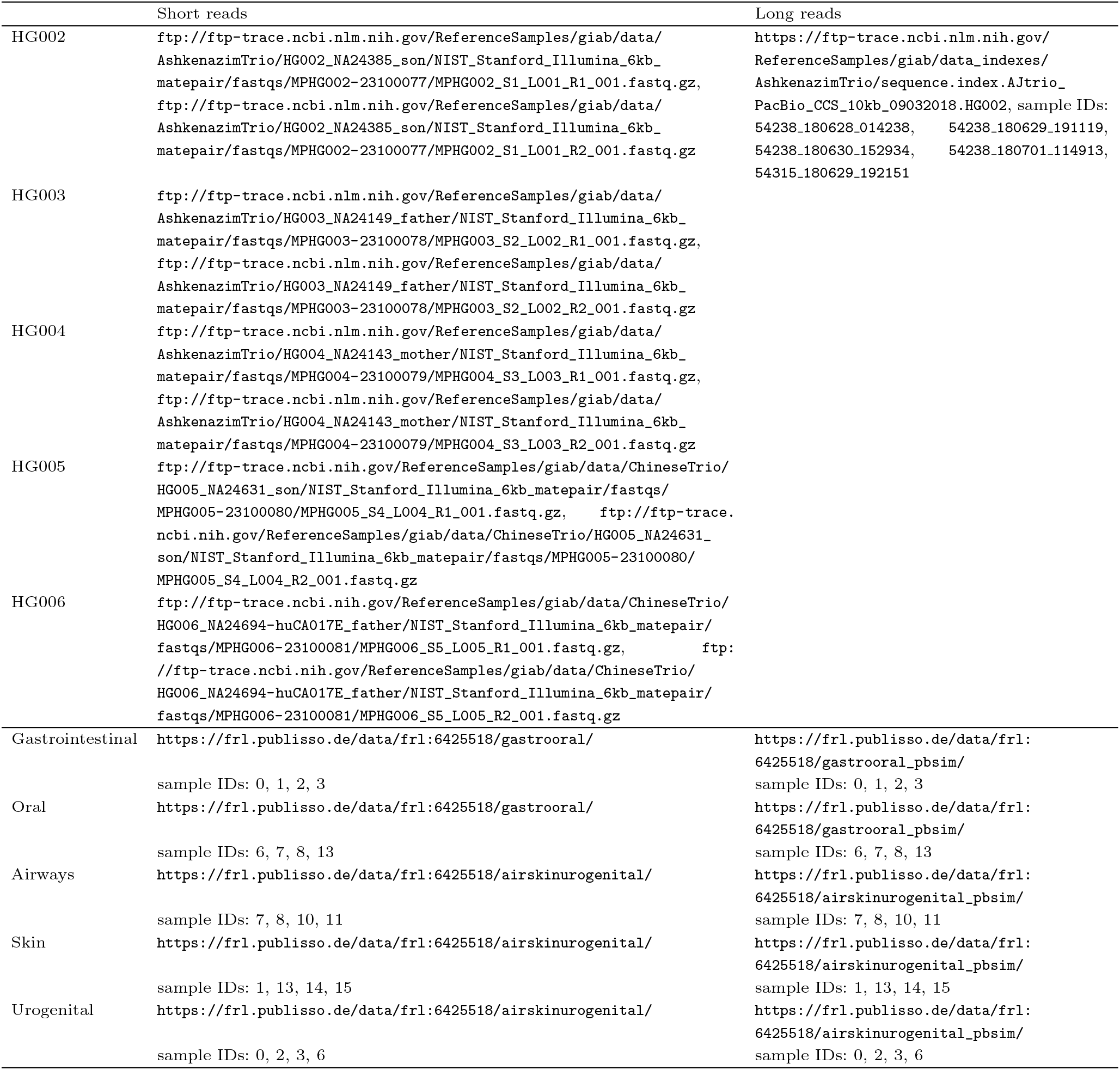
Data URLs for human (GIAB) and microbiome (CAMI 2 challenge) datasets. The data is subsampled (for human) or combined (for microbiome) such that each file contains 50 million short reads or 5 million long reads.

## Conclusion and Discussion

We presented a new tool, called Cleanifier, for removing host contamination from microbiome sequencing projects. Cleanifier is very fast in sampling mode, relatively memory-efficient (it will run on an 8 GB system using the probabilistic Cuckoo filters) and offers consistently high accuracy on both short and long read data. In addition, Cleanifier supports building custom databases for other host organisms, e.g., for removing mouse reads from mouse gut microbiome samples.

The comparison between Cleanifier using a Cuckoo hash table (exact) and using Cuckoo filter (probabilistic) shows that under the same circumstances, i.e., using the same references to build the index and applying the same classification method, the small FPR incurred by the probabilistic Cuckoo filter does not negatively affect the accuracy on short or long reads. However, results on very short reads (such as 50 bp reads sometimes used for transcript quantification) may be different, if a single gapped *k*-mer hit might decide the classification.

To improve Cleanifier further, future work could address new sampling schemes, such as more advanced skipping heuristics, or using minimizers (Roberts et al., 2004) or syncmers (Edgar, 2021) for sampling. Furthermore, a new human index may consider masking genomic regions that are known to have a high similarity with microbial sequences to enhance microbiome retention.

## Code Availability

The code and Snakemake workflows (Mölder et al., 2021) to reproduce all results are available via GitLab (https://gitlab.com/rahmannlab/cleanifier). In addition, Cleanifier is available at PyPi (https://pypi.org/project/cleanifier/) and Bioconda (https://anaconda.org/bioconda/cleanifier).

## Data Availability

All data used in the evaluation is available online (Meyer et al., 2025; Zook et al., 2016). Dataset URLs are provided in the Supplement. The pre-built Cleanifier index is available at Zenodo (https://doi.org/10.5281/zenodo.15639519) or can be downloaded using the `cleanifier download` command.

## Supplementary Material

### Dataset URLs

### Cleanifier Speedup Factors

**Fig. S.1:**
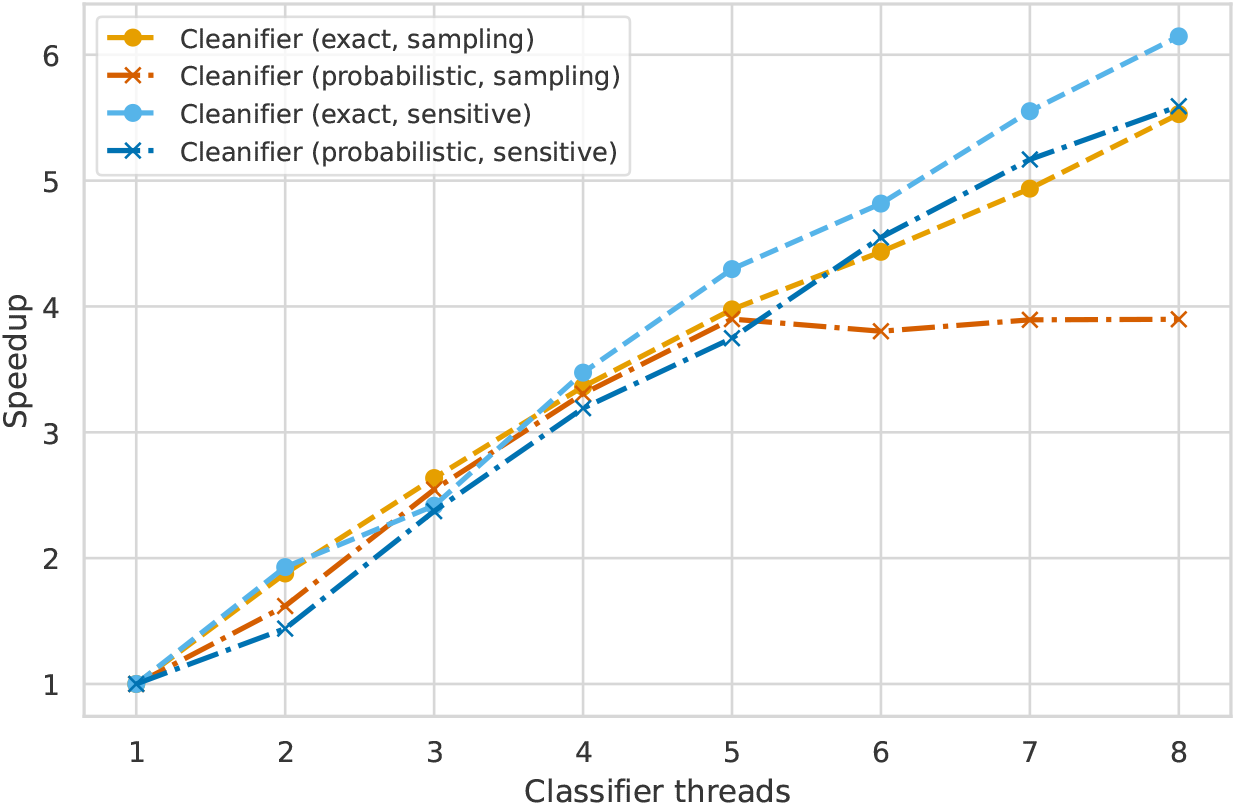
Speedup for increasing classifier threads. Total threads are the number of classifier threads plus one additional read and one additional write thread.

The speedup factor from parallelization is almost linear for up to 8 classifier threads except for the probabilistic sampling version (Figure S.1). Here, the read or write thread becomes the main bottleneck if more than 5 threads perform the classification.

### Accuracy for different classification thresholds and *k*-mer masks

We built indexes for 6 different *k*-mer masks that were selected from the best performing *k*-mer masks evaluated in Rahmann and Zentgraf (2023), see Table 4. Figure S.2 shows the accuracy for human and microbiome data for the different masks and different classification thresholds. Due to the good performance on both human and microbiome data, we selected the (29,33)-mask and set the default threshold for classification to 0.5.

**Table 4.**
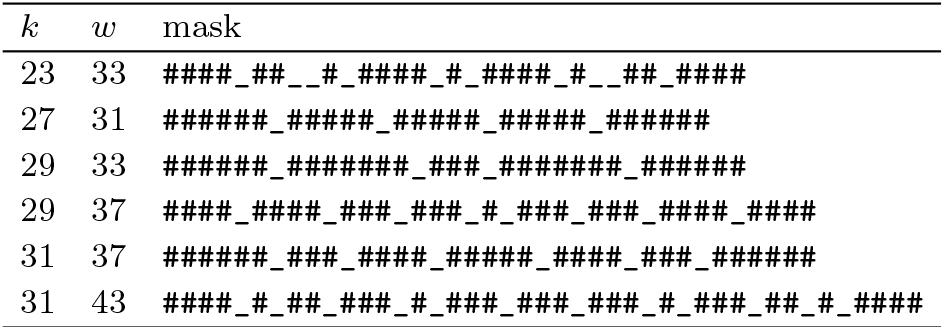
Evaluated gapped *k*-mer masks.

**Fig. S.2:**
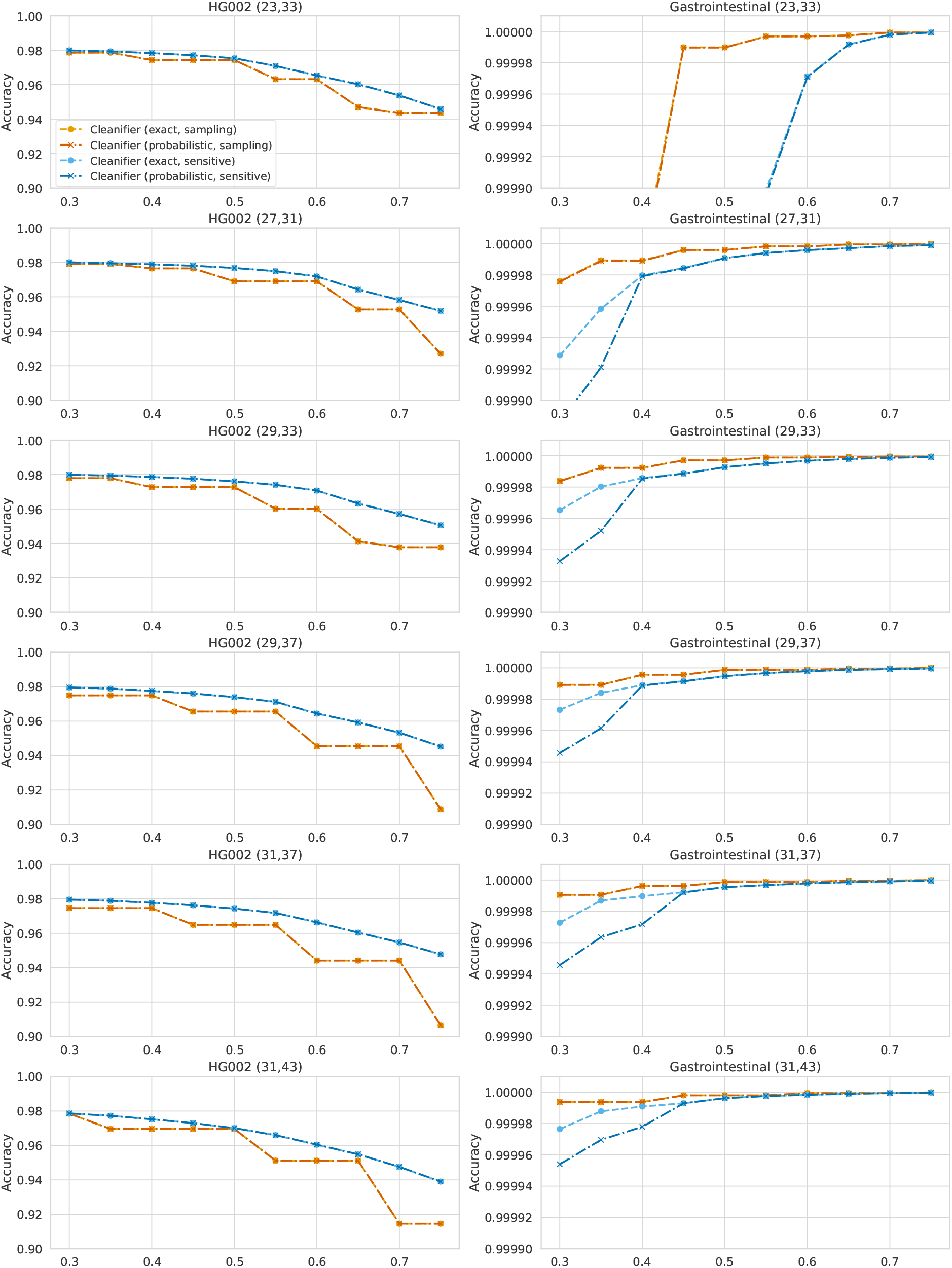
Accuracy for human and microbiome data for different indexes (built using different gapped *k*-mer masks) and different classification thresholds.

**Fig. S.3:**
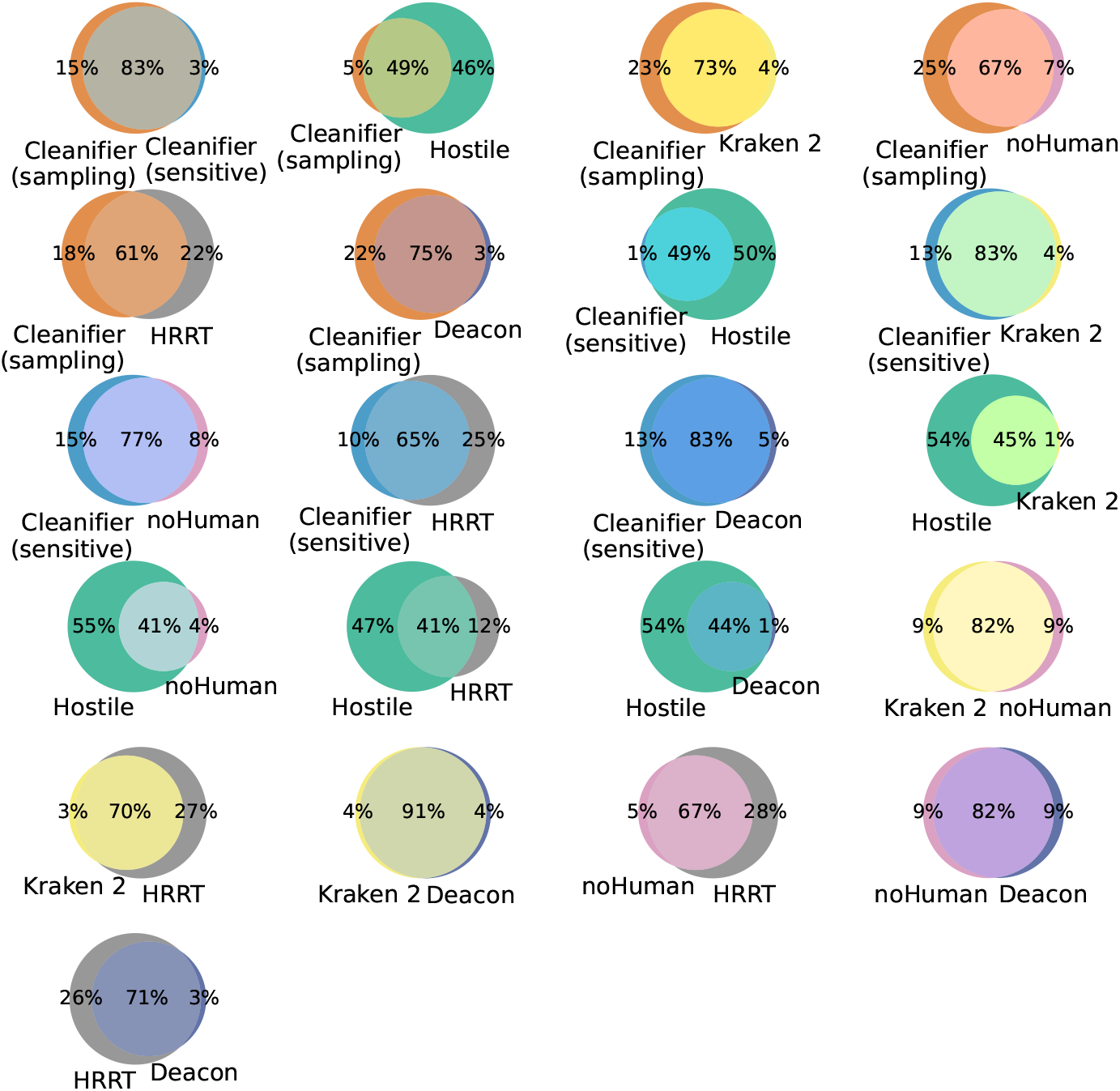
Overlap of retained reads of the HG002 dataset between all tools. We only included Cleanifier (probabilisitc) in the comparison.

### Comparison of T2T and the advanced human index

Table 5 shows the accuracy for an index containing only the gapped *k*-mers from the T2T reference Nurk et al. (2022) and from our advanced human index that is built from the T2T reference, the common variants from the 1000 Genome Project Auton et al. (2015), the HLA variants from the IPD-IMGT/HLA database Barker et al. (2023) and the 47 assemblies from the Human Pangenome Consoritium Liao et al. (2023). Our advanced index leads to a higher accuracy on human data both for the exact and probabilistic version, with a higher increase in the accuracy for the sampling mode. The microbiome retention is only marginally reduced by the more comprehensive human pangenome index.

**Table 5.** Accuracy for an index built for the T2T reference and for an index containing all gapped *k*-mers of the T2T reference, common variants, HLA variants and multiple pangenome assemblies. We measured the accuracy on the HG002 and gastrointestinal short read datasets for a threshold of 0.5.

|  | Cleanifier exact |  |  |  | Cleanifier probabilistic |  |  |  |
| --- | --- | --- | --- | --- | --- | --- | --- | --- |
|  | sampling |  | sensitive |  | sampling |  | sensitive |  |
|  | T2T | pangenome | T2T | pangenome | T2T | pangenome | T2T | pangenome |
| Human | 0.969986 | 0.972677 | 0.975124 | 0.976031 | 0.969988 | 0.972680 | 0.975136 | 0.976045 |
| Microbiome | 0.999999 | 0.999997 | 0.999996 | 0.999993 | 0.999999 | 0.999997 | 0.999996 | 0.999993 |

### Pairwise overlap between retained human reads

Figure S.3 shows the pairwise overlap of the sets of reads that retained in the HG002 dataset by all evaluated tools. Since all tools have a high overlap, it is possible that many of these reads are either contaminants, like PhiX or the Eppstein-Barr virus, or poor quality reads and should indeed be retained as non-human.

